# Experimental evolution reveals competing selective forces during an early pollinator shift in monkeyflowers

**DOI:** 10.64898/2026.09.28.755135

**Authors:** Foen Peng, Claudia van Vloten, Benjamin Winograd, Abigail Trapp, Sophia Taberski, Lexie Streicher

## Abstract

Understanding how complex multi-trait adaptations arise in nature remains a fundamental challenge in evolutionary biology. A classic example of this adaptation is the pollination syndrome, where integrated suites of floral characteristics co-evolve to attract specific pollinators. While transitions between pollination syndromes occur frequently, the underlying evolutionary trajectories traversed by adapting populations remain largely opaque. Here, we used an experimental evolution approach to track real-time phenotypic and genomic changes during a pollinator switch, leveraging the novel attraction to bumblebees exhibited by hybrids between the hummingbird-pollinated *Mimulus cardinalis* and the self-pollinating *M. parishii*. Within only three generations of bumblebee selection, we observed remarkably rapid shifts in floral traits, genotypes, and fitness. Our findings demonstrate that pollinator shifts are likely shaped by a complex interplay of selection, genomic conflict, and pleiotropic constraints that cannot be predicted simply by floral traits’ genetic architecture alone.

## Introduction

Adaptation often involves concerted shifts across multiple traits. Prominent examples span diverse systems, from cavefish adapting to dark, nutrient-poor environments via eye loss, enhanced chemosensory capabilities, and improved metabolic efficiency, to domesticated crops exhibiting a “domestication syndrome” characterized by reduced seed shattering, higher yields, and decreased branching. In flowering plants, a classic example of multi-trait adaptation is the pollination syndrome, where integrated suites of floral traits, such as color, scent, and morphology. co-evolve to attract specific functional groups of pollinators (Fenster et al., 2004). Transitions between distinct pollination syndromes occur frequently in nature; for example, transitions between bee and hummingbird pollination have occurred over 100 times in North America alone (Thomson & Wilson, 2008). Yet, despite extensive documentation of this phenomenon, how these genetically complex adaptive syndromes arise repeatedly, and often rapidly, remains a fundamental mystery in evolutionary biology.

When distinct pollination syndromes are viewed as discrete peaks on a fitness landscape, several non-mutually exclusive hypotheses explain how plant populations cross the intervening fitness valleys. Adaptive walk models suggest that populations typically navigate toward new fitness peaks through a series of beneficial mutations characterized by diminishing returns (Orr, 1999). Under this framework, initial large-effect mutations bridge fitness valleys by shifting a population near a new phenotypic optimum, while subsequent small-effect mutations cumulatively fine-tune the adaptation. In the context of pollination syndromes, this model predicts that pollinator attraction traits (like color and scent) and reward traits (like nectar volume) should evolve early during a transition, whereas dimensional traits (like corolla length) are optimized later (Wessinger, 2024), because the genetic architecture of dimensional traits tends to be more complex (Cui & Yuan, 2026). Alternative frameworks invoke mechanisms such as delayed selfing (Wessinger & Kelly, 2018), secondary pollinators (Kay & Anderson, 2025), or hybridization (Rezende et al., 2020) to stabilize populations during intermediate stages of a transition. However, direct experimental data testing these hypotheses remain scarce.

Experimental evolution offers a powerful approach to illuminate these otherwise opaque evolutionary trajectories. For example, domesticated animals frequently exhibit a “domestication syndrome”—including docile behavior, reduced skull size, and floppy ears—that puzzled Darwin himself (Wilkins et al., 2014). To resolve this, Russian scientists in the 1960s initiated an experiment to domesticate silver foxes. Within just a few generations, foxes selected strictly for tameness began displaying the morphological traits of domestication syndrome, demonstrating that these physical changes were pleiotropic byproducts of selection on behavior (Trut, 1999). In plant systems, experimental evolution has similarly yielded critical insights. For instance, recent work in *Arabidopsis* demonstrated that rapid climate adaptation can occur within a few generations (Wu et al., 2026). Furthermore, *Mimulus guttatus* evolved self-pollination in as few as five generations following pollinator loss (Bodbyl Roels & Kelly, 2011), and *Brassica rapa* floral traits diverged rapidly under reduced pollinator diversity (Gervasi & Schiestl, 2017). Nevertheless, no experimental studies have tracked how a plant population evolves to adapt to an entirely novel pollinator group.

Recently, we identified a unique opportunity to tackle this challenge. We found that the F1 hybrids between the hummingbird-pollinated *Mimulus cardinalis* and the self-pollinating *Mimulus parishii* unexpectedly attract bumblebees, a pollinator visiting neither progenitor species (Peng et al., 2024). Despite their close phylogenetic relationship, *M. cardinalis* and *M. parishii* have distinct pollination syndromes (Fig. 1A). *M. cardinalis* exhibit large corollas, vivid red pigmentation, copious nectar rewards, and exerted reproductive organs, while *M. parishii* has miniature-sized flowers with no nectar and has minimal anther-stigma separation. Advanced generations of these hybrids segregate for a myriad of floral traits without experiencing notable inbreeding depression (Liang et al., 2023). Given the novel attraction to bees, experimentally evolving the hybrid population under bumblebee selection would enable a detailed study on the phenotypic and genomic changes during a pollinator syndrome switch. It is worth to note that within the same clade of *Mimulus*, there is another closely-related species *Mimulus lewisii*, which is naturally pollinated by bumblebees and has a typical bumblebee pollination syndrome: pink corolla, small amount of concentrated nectar, inserted reproductive organs, and distinct landing platforms. Comparing the experimentally evolved hybrid lines with the naturally evolved *M. lewisii* could reveal whether similar evolutionary paths were repeatedly taken during adaptation to the same pollinator.

**Figure 1.**
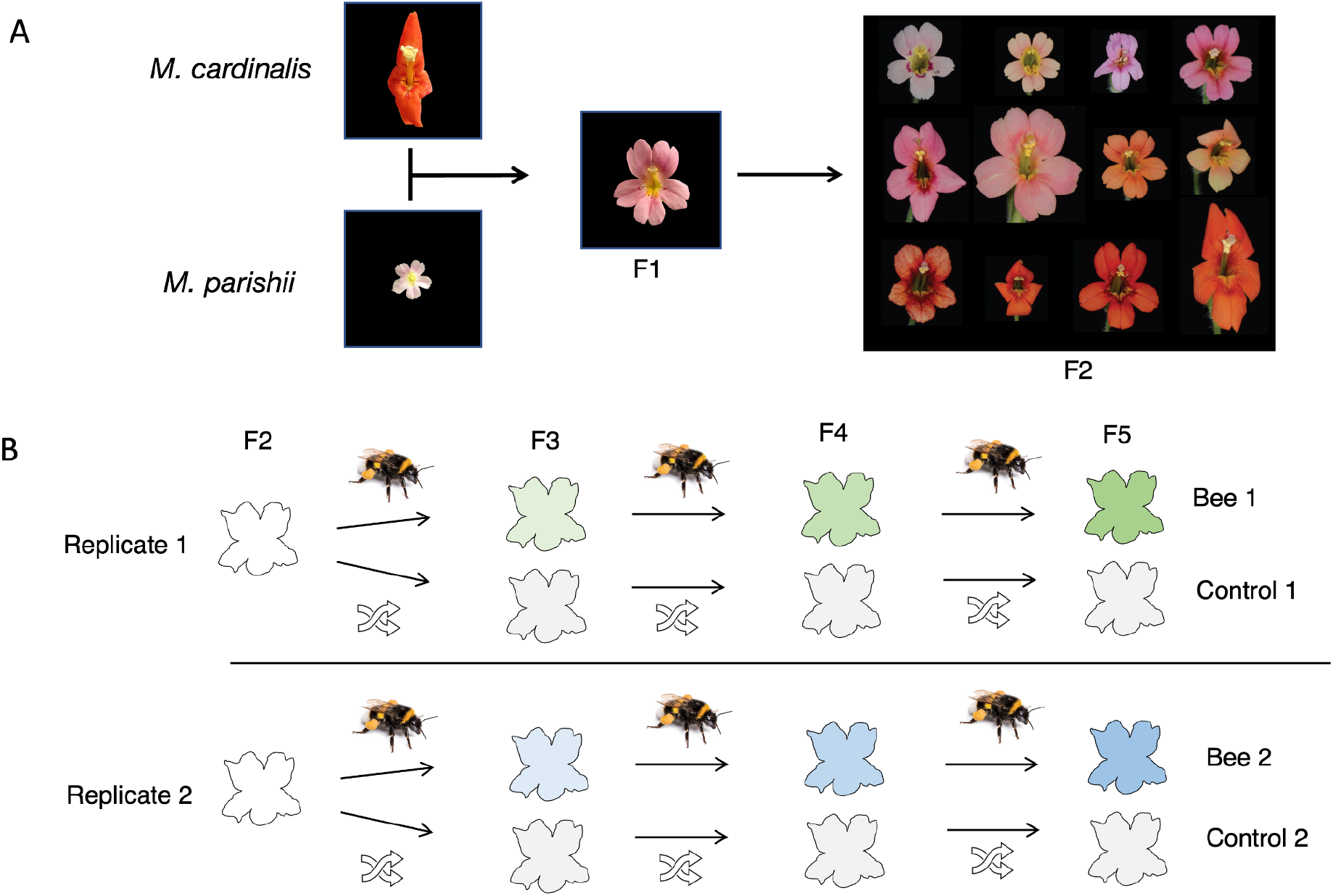
A. The patentai species. F1. and F2 hybrids flowers. B. The evolution experiment design

## Materials and Methods

### Plant materials

The *M. cardinalis* inbred line CE10 and the *M. parishii* inbred line PAR were used to establish the hybrid population for the evolution experiment. The seeds of both inbred lines were obtained from the Yuan Lab at University of Connecticut, and the hybrid seeds were generated at Haverford College, PA. The F1 hybrids were generated by a cross using *M. cardinalis* CE10 as the pollen donor and *M. parishii* PAR as the pollen receiver. The F1s were self-pollinated to generate the F2 seeds. Plants were grown in the Haverford College greenhouse under long day conditions (16hr light / 8hr dark). They were watered daily with a dilute fertilizer solution.

### Floral traits measurement

Newly opened flowers were labeled every day during the measurement period and only flowers that were 1-2 days old were used in trait measurements. A total of 15 traits in were measured for each plant in every generation, including corolla tube length (CTL), corolla tube width (CTW), corolla lobe length (CLL), corolla lobe width (CLW), the length of the longer set of stamens (stm_len), pistil length (pst_len), stalk length (stk_len), nectar volume (nec_vol), nectar concentration (nec_con), projected area (prj_area), petal color (R, G, B), petal reflex (reflex) and nectar guide spot size (dot_size). Flower volatiles were not collected because neither parent species (Peng et al., 2017) or the F1s (unpublished data) produce bee attracting volatiles. We used a digital caliper for all the shape measurements. We selected two recently opened flowers per plant for shape measurements. For example, corolla tube length (CTL) was measured as the distance from the receptacle to the petal split between dorsal and lateral petals. For a detailed explanation of all the shape variables, see (Chen, 2022). Nectar volume and nectar concentration were also recorded from two flowers per plant. Nectar volume was measured with 25uL Microcaps disposable glass micropipettes (Drummond) and nectar sugar concentration was measured with an Eclipse low volume refractometer. The petal reflex was visually scored on 3-5 flowers per plant as complete reflex (similar to *M. cardinalis*), intermediate reflex, or no reflex (similar to *M. parishii*). Similarly, nectar guide spot size was visually scored as large connected areas, intermediate, or small unconnected dots. The front view digital photo of one flower for each plant was recorded by an Olympus TG-6 camera in microscope mode (no flash) in a dark room in ambient room light at a fixed distance from the flowers.

To accurately record flower color, we took reflectance spectra for one flower of each plant, starting from F3. We also took reflectance spectra for flowers from *M. cardinalis, M. parishii, M. lewisii*, the F1 between *M. cardinalis* and *M. parishii*, as well as a representative leaf sample. For each flower, the petal lobe area was illuminated using a PX-2 pulsed Xenon light source, triggered by a 200Hz 5V TTL function generator pulse. An Ocean Optics USB2000+RAD spectrophotometer was used for data collection with OceanView software parameters: reflectance mode, 30ms integration time, averaging 5 consecutive scans, and a pixel boxcar average of 3. We used a WS-1 Diffuse Reflectance Standard (OceanInsight) for the white standard and the complete darkness in a dark room to define dark counts. Reflectance spectra were processed using the R packages lightr and pavo (Maia et al., 2013). We used the color hexagon model (Skorupski & Chittka, 2010) to model bee vision as it is specifically constructed for hymenopteran vision. We used 347, 424 and 539 nm for UV, blue and green receptors to model the color hexagon model (Skorupski & Chittka, 2010).

### Evolution experiment design

F2 hybrids were used as the founding populations. An initial F2 population was first subject to bumblebee pollination to start one experimental line, then randomly hand-pollinated to start a control line, which evolved in parallel with the experimental line (Fig. 1B). The replicate experimental and control lines started from a distinct F2 population and underwent the same procedures. As a result, starting from F3, there were two experimental populations and two control populations evolved in parallel. Each population constituted 50 individuals and the experiment continued for 3 generations, with one generation in each year of 2023, 2024, and 2026.

### Bee pollination procedures for experimental populations

Each experimental population was subject to bumblebee pollination for 4 days. The bumblebees were artificially reared flower-naive *Bombus impatiens* from a commercial source (Plant Products’ BioBest standard hives). The hives included a queen and over 100 workers. The hives had access to sugar water and were fed with fresh pollen (also from Plant Products) every other day. The hives were acclimated to a flight cage (Growneer crop cage, Amazon), which was made of netting materials with a dimension of 3m (L) x 2m (W) x 2m (H), for one week before the pollinator visitation experiment. The flight cage was put outside of the Haverford College greenhouse so that the bees foraged on flowers exposed to natural sunlight. During acclimation of the bees, 30-40 plants from the hybrid populations each day were placed in the cage, none of which were used in the actual experiment later. Sugar water supply to the hives was cut off 16 hours before each experiment day to motivate the bees to search for food; this did not result in a noticeable negative impact on hive health. During the experiment, the bees in the hive were allowed to forage freely within the flight cage and return to the hive. The 50 hybrid plants in the cage were placed in a 6 x 9 array and the positions were randomized at the beginning of every experiment day. Two F1 plants, with all flowers emasculated to avoid pollen contamination, were also placed in the flight cage as control. The number of flowers on each plant was counted before the start of the experiment. The experiments were conducted in late April to early May each year. Typically, there were 5-15 bees foraging in the cage at any given point. The experiment was observed by two observers for multiple 20-min windows between 1pm and 4pm. During those periods, the observers recorded the number of visits to each specific plant. A visit was counted when a bee made physical contact with the anther or stigma of a flower, or the bee crawled into the corolla tube to access nectar. At the end of each experiment day, all the floral corollas are removed, so that only newly opened flowers are available for bees to forage in the next experiment trial. To avoid seeds generated by selfing pollen in experimental lines, two flowers from each plant were emasculated prior to exposure to bees and subsequently used for seed collection; other flowers on the same plants were not emasculated to serve as pollen donors.

### Random hand pollination procedures for control populations

For each control population, a random number series was provided by the *sample* function in R 4.3.1 (R Core Team, 2021). For any given plant in the control, two flowers were pollinated with pollen from the previous plant in the random series. Then, the focal plant served as the pollen donor for the next plant in the series. As a result, each plant in the population only received pollen from one plant and provided pollen to pollinate another plant. During hand pollination, several anthers of a flower were rubbed against the recipient flowers’ stigma to ensure sufficient pollination. This procedure ensured that there was no selection in the pollination process.

### Seed harvesting and pooling

Seeds were harvested 3-4 weeks after pollination. Fruits from the same plants were pooled together for seeds counting. We performed seeds counting with a custom computer vision script in ImageJ based on background subtraction. For the experimental populations, the number of seeds per fruit was used as a fitness metric, and the percentage of seeds each plant contributed to the next generation was proportional to their relative fitness in seeds production. For each of the control population, 10 seeds from each plant were pooled to produce the seed bank for the control population of the next generation.

### Genotyping

The DNA of each individual was extracted from 3-5 young buds using NucleoSpin Plant II, Mini kit (Macherey-Nagel). A pair of primers was designed to perform PCR-based genotyping for the *YUP* locus (Forward: TGGGTTGGTTATGGTCGATT; Reverse: CGTGAAAGCTTTACACATGTCT) by amplifying a region adjacent to the YUP siRNA gene (~2kb upstream). The amplicon from *M. parishii* allele was 312 bp long while the amplicon from *M. cardinalis* allele was 470 bp long. We used GoTaq Master mix (Promega) for the PCR reactions. The following PCR protocol was used: step 1, 95°C for 3 min; step 2, 95°C for 30 s; step 3, 55°C for 30 sec; step 4, 72°C for 30 sec; 40 additional cycles of steps 2 to 4; finish at 72°C for 5 min. The PCR product was stained with EtBr and run on a 1.4% agarose gel to visualize.

### Statistical analysis

Data analysis was conducted with R 4.3.1(R Core Team, 2021). In order to calculate the bumblebee visitation rate for each plant, which was defined as visits per flower per hour, we divided the total number of visits over all experiment days by the total number of flowers and the total number of observation hours.

## Results

### 1. Experimental populations increased fitness under bee pollination

Plants in the experimental populations became attractive to bees over time (Fig. 2). For this analysis, we used F1s as a baseline reference due to their phenotypic uniformity and moderate attractiveness to bees. Within the initial F2 populations, the majority of individuals were less attractive than the F1 reference; only 16% and 12% of the plants across the two replicates received more pollinator visits than the F1 baseline. However, after three generations of bee-mediated selection, the proportion of plants outperforming the hybrids climbed to 40% and 30% in the respective F5 experimental populations. This directional trend was even more pronounced at the upper tail of the phenotypic distribution. While the most attractive plants in the F2 replicates received only 49% and 19% more visits than the F1 control, the most attractive individuals in the F5 generation attracted 240% and 184% more visits than the F1 hybrids, respectively.

**Figure 2.**
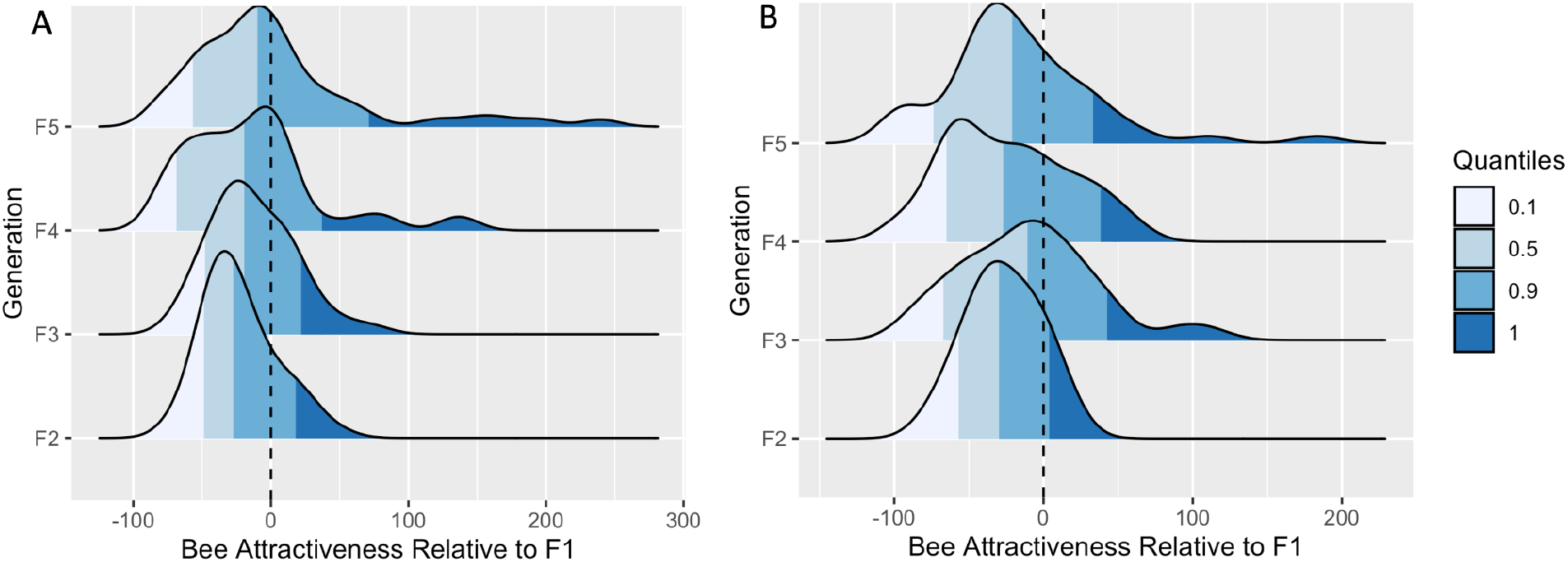
A. and B. The distribution of bumblebee attractiveness in the two experimental populations, respectively. Attractiveness was calculated as the total number of visits per flower per hour, normalized by F1 attractiveness.

### 2. Rapid phenotype changes were observed in many traits

Many floral traits showed a high degree of correlation (Fig. 3A). They formed three clusters on the correlation plot. Floral color, measured as the amount of reflectance for petals in the three peak sensitive wavelengths, are correlated with each other. The two relative length measures, sigma-anther separation and stigma to corolla distance, were correlated with each other. All other shape traits, along with nectar volume form the third major cluster. Petal dot size, nectar concentration, and stigma closing speed did not show a strong correlation with other traits.

**Figure 3.**
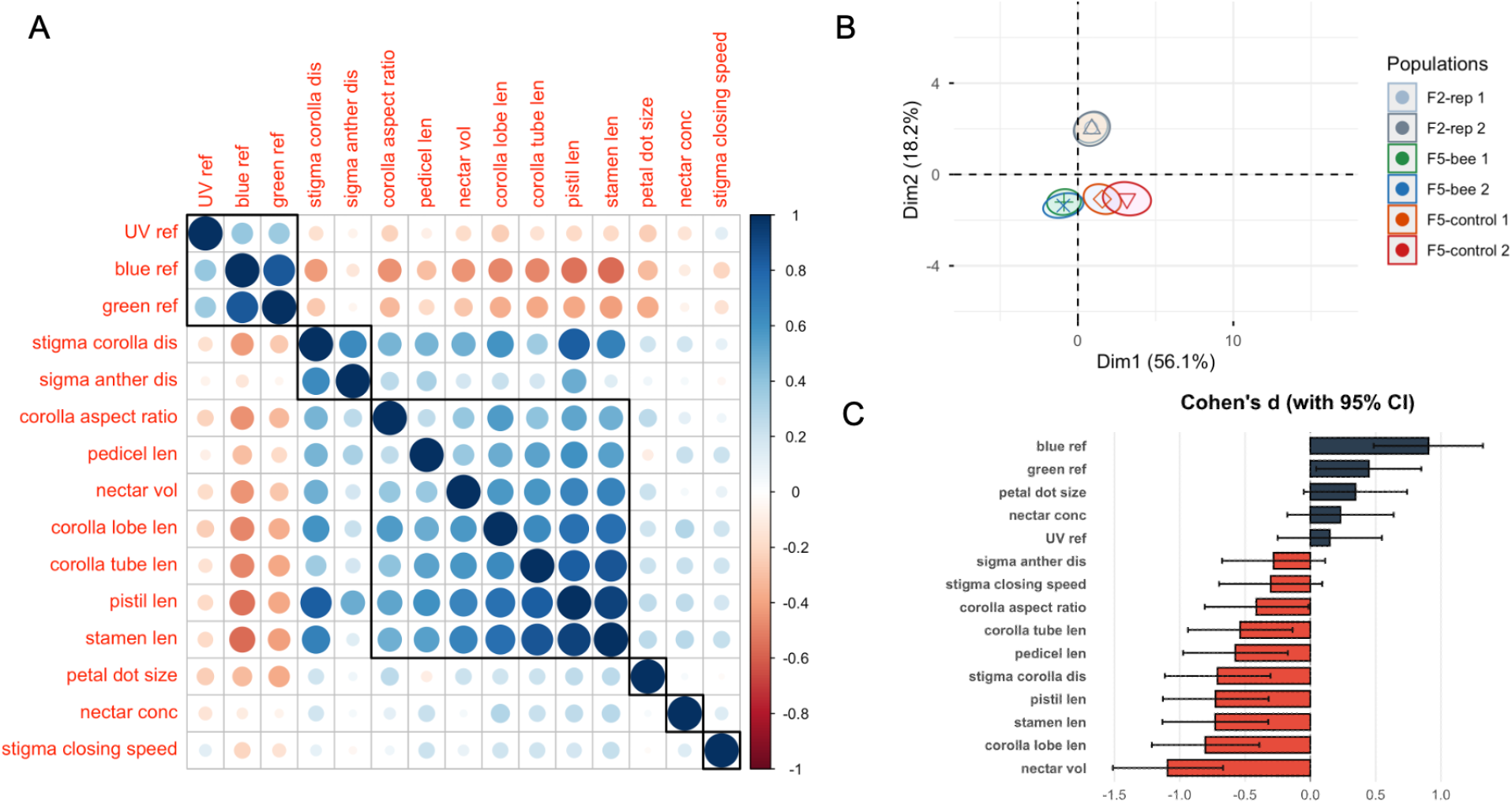
A. Correlation plot for all the floral traits. B PCA for all the traits in F2 and F5 populations. C. Average trait divergence between experimental and control populations in F5. measured as Cohen’d.

When plotting the floral traits on PCA space, the first two principal components (PC1 and PC2) accounted for 74.3% of the total floral phenotypic variation (Fig. 3B). The two replicate F2 populations overlapped almost completely, indicating minimal variation between the randomly sampled founding groups. By the F5 generation, the two bee-pollinated populations had diverged sharply from both the ancestral F2 and the F5 controls. Despite this divergence, they clustered closely with one another. Intriguingly, the two F5 control replicates exhibited divergence from the ancestral F2 population as well as moderate divergence from one another, suggesting that other processes, such as genetic drift or unintended selection, may have also shaped these populations.

When examined individually, many traits showed significant divergence between the experimental and control populations in F5 (Fig. 3C). Specifically, petal color had stronger blue and green reflectance in the experimental populations. Nectar volume was significantly reduced in the experimental populations, and almost all the shape traits became smaller or shorter.

### 3. Experimental populations had significant phenotypic changes in many floral traits

Previous work (Peng et al., 2024) showed that bumblebees preferred flowers with large corolla, therefore, we did not expect the experimental lines to evolve an overall smaller flower. To understand the mechanisms behind this trend, we examined one particular shape trait, pistil length, in more detail. While the control populations had similar pistil length through the generations, the experimental populations had shorter and shorter pistils (Fig. 4A). We found that pistils had no impact on bumblebee attractiveness (t.test, p=0.28) (Fig. 4B), but the longer pistil significantly reduced the seed set (t.test, p<0.001) (Fig. 4C).

**Figure 4.**
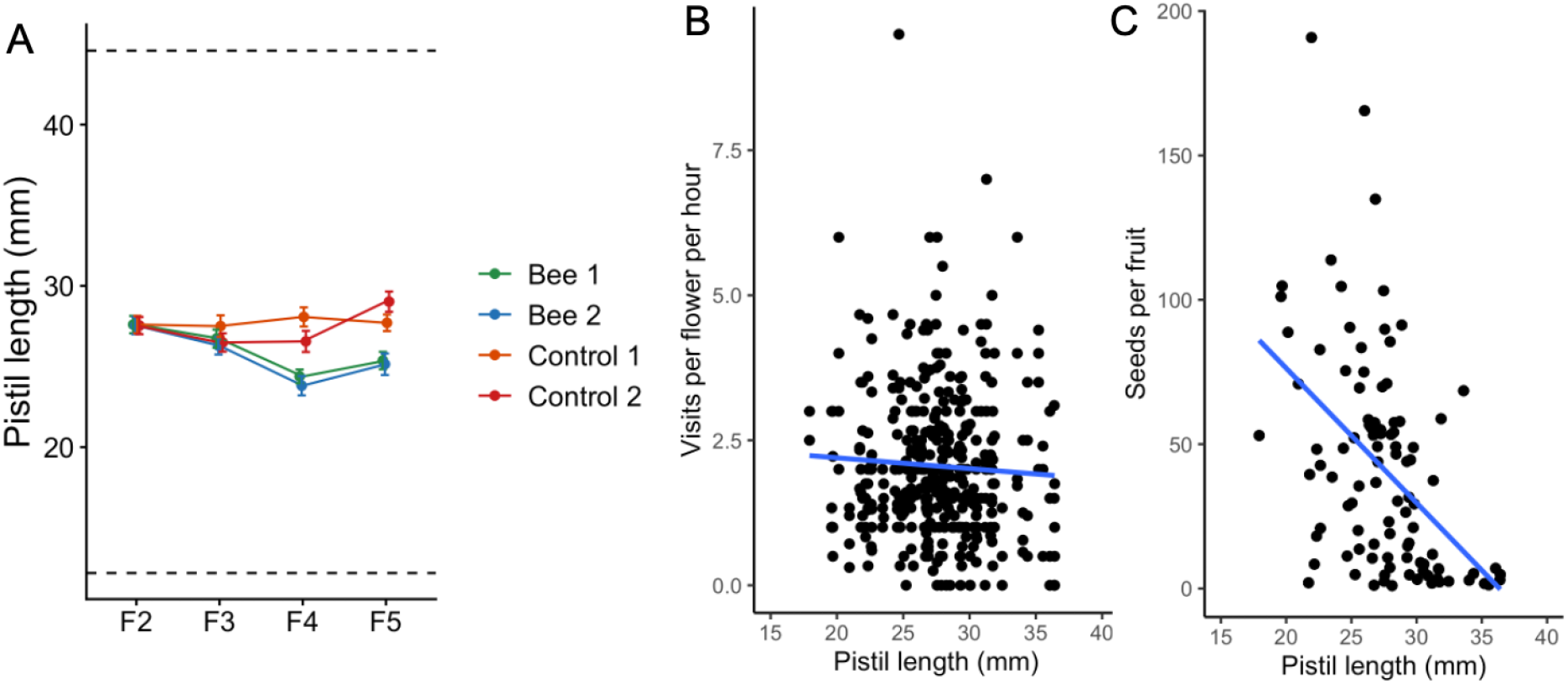
A. Change of pistil length over time. B. Regression of pistil length with visitation rate. C. Regression of pistil length with average seed set per fruit.

### 4. Divergence of allele frequencies between experimental and control populations

Given the pronounced evolutionary shifts in petal color, we expected corresponding allele frequency changes at the underlying genetic loci. A primary candidate is *YUP*, a locus encoding a small RNA gene that represses carotenoid synthesis; the dominant *M. parishii* allele (P) leads to pink coloration, accounting for much of the color divergence between the parental species. Consistent with the observation that bee-pollinated flowers evolved increased reflectance in the blue wavelength spectrum (Fig. 5A), the P allele frequency steadily rose in these populations (Fig. 5B).

**Figure 5.**
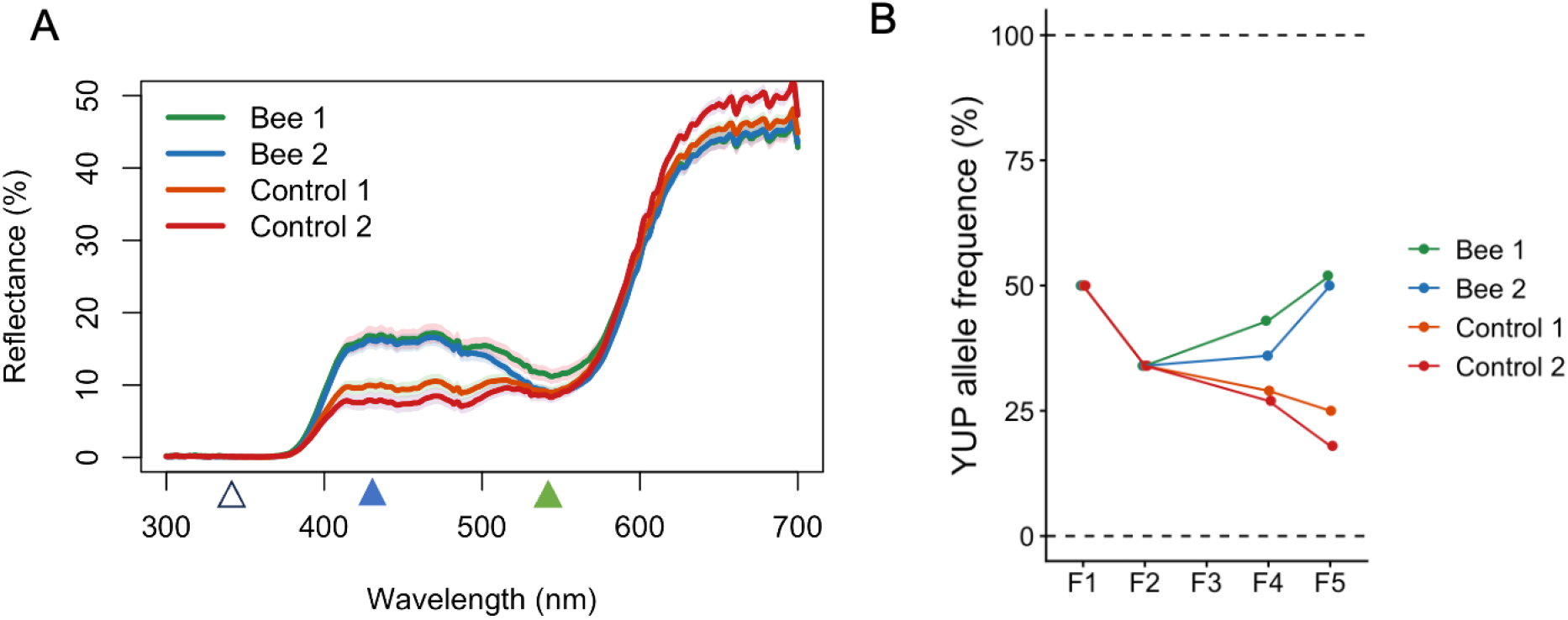
A. Petal reflectance spectra for F5 populations. Three sensitive peaks of bee’s color cones were indicated as triangles. B Change of *M parishii* YUP allele frequency over time.

Intriguingly, the initial F2 populations both exhibited a P allele frequency of only 34%, deviating significantly from the Mendelian expectation of 50% under free segregation (Chi-squared test, p < 0.001). This pattern suggests segregation distortion operating against the transmission of the P allele. In the absence of counteracting selection, such distortion should drive a continuous decline in allele frequency over subsequent generations. This expectation was confirmed in the two F5 control populations, where the P allele frequency dropped to 25% and 18%, respectively.

Although segregation distortion must have similarly penalized the P allele transmission in the bee-pollinated lineages, directional selection mediated by pollinators was powerful enough to overcome this transmission deficit. Consequently, the P allele frequency surged to 52% and 50% in the experimental replicates. This tug-of-war between segregation distortion and pollinator preference generated substantial divergence between the experimental and control populations, resulting in a net allele frequency difference of 27% and 32% across the two replicates.

## Discussions

Within only three generations of evolution experiment, we observed remarkably rapid shifts in genotype, phenotype, and fitness. Multiple floral traits, including color, shape, and rewards, responded to selection imposed by the new bumblebee pollinator. Although evolution was traditionally viewed as a slow and gradual process, rapid evolution has been frequently documented over recent decades (reviewed in Hairston Jr et al., 2005). Consistent with other experimental evolution studies in plant systems (Bodbyl Roels & Kelly, 2011; Gervasi & Schiestl, 2017; Wu et al., 2026), our findings demonstrate that selection-driven evolutionary changes can occur rapidly, even within small populations where genetic drift is conventionally expected to be a dominant force.

Floral traits can be categorized based on their distinct ecological functions (Bradshaw et al., 1998): pollinator attraction traits (e.g., color and scent), pollinator reward traits (e.g., nectar volume and concentration), and pollination efficiency traits (e.g., pistil length and petal size).

Among the traits measured in this study, color and shape traits formed distinct clusters in our correlation analysis. This within-cluster correlation suggests that selection acting on any single trait is likely to drive correlated evolutionary changes in others. This pattern of phenotypic integration has been widely reported in other plant systems (Smith, 2016), establishing the genetic backdrop against which selection operates. Interestingly, we also noticed some exceptions to this general clustering pattern. We found that petal lobe color was uncoupled from the size of the petal dots within nectar guide regions. This independence aligns with the fact that distinct MYB transcription factors regulate coloration in the petal lobe versus the nectar guide zones (Yuan et al., 2014). Furthermore, while nectar volume was highly correlated with multiple floral shape traits, nectar concentration showed no such association. This divergence suggests that these two reward traits rely on distinct or unlinked genetic bases. Because pollinators frequently rely on flower size as an honest signal to gauge reward quantity (Parachnowitsch et al., 2019), the genetic coupling of nectar volume with floral morphology may provide an adaptive advantage.

Previous work found that pollinator attraction traits and pollinator rewarding traits tend to be controlled by a few large effect loci, while pollination efficiency traits are more likely to be controlled by many loci with small effect (Cui & Yuan, 2026). This genetic architecture predicts a distinct evolutionary sequence: attraction and reward traits should evolve fast during a peak shift, whereas dimensional fine-tuning occurs later (Wessinger, 2024). In contrast with this prediction, we found that all three types of traits, including petal color, floral shape, and nectar volume, evolved rapidly in our experiment, which suggests that the genetic bases alone is not enough to predict the evolutionary rate of floral traits in pollinator shift evolution.

Instead, the results from this early phase reveal a highly dynamic landscape wherein several selective forces jointly shape evolutionary trajectories. For example, the shift in petal color is primarily driven by a strong preference of bumblebees for pink over orange-red coloration (Peng et al., 2024). Because pink reflects more strongly in the blue region of the light spectrum, it is presumably more conspicuous to bees. Previous research established that *YUP*, a Mendelian locus controlling carotenoid pigmentation, is largely responsible for the color variation in this cross (Liang et al., 2023). Interestingly, by analyzing *YUP* allele frequencies over evolutionary time, we discovered that strong pollinator-mediated selection on floral color was likely counteracted by a segregation distortion locus acting in the opposite direction (Sotola et al., 2023). Yet, despite this tug-of-war between segregation distortion and bee selection, substantial color divergence emerged between the experimental and control populations after only three generations. Regarding dimensional traits, several traits evolved to be smaller or shorter. For instance, pistils became significantly shorter in bee-pollinated F5 individuals. This shift is explained by our observation that a shorter pistil facilitates more effective pollen deposition, thereby increasing seed set. However, pollination efficiency alone cannot account for the reductions observed in other dimensional traits, such as corolla lobe, pedicel, and corolla tube lengths. Given our previous finding that bees prefer visiting flowers with larger corollas, one would predict the evolution of larger, rather than smaller, floral structures. We speculate that this general reduction in flower size stems from a tight phenotypic correlation between pistil length and other dimensional traits; in this case, the selective pressure optimizing the quality of pollinator visits overbalanced the preference for quantity. A similar mechanism likely explains the observed reduction in nectar volume, given its tight correlation to other dimensional traits.

Although the evolution experiment was designed to minimize confounding factors, it revealed a highly complex landscape of multiple forces acting jointly. These include the action of selfish genetic elements (*i*.*e*., segregation distortion) and the conflicting selection pressures balancing pollination quality against quantity. As a result, the experiment has provided unique insights into the real-time dynamics of pollinator shifts that is hard to be inferred retrospectively from extant wild populations. Moving forward, tracking the long-term evolutionary trajectories of these populations will be interesting. For instance, will strong bee selection lead to the fixation of *YUP* alleles in the experimental populations, while control populations fix for the alternative allele? Furthermore, once the pistil becomes short enough to ensure successful pollination, will bee preference drive the evolution of larger flower sizes, given that the bee-pollinated relative *Mimulus lewisii* exhibits both a short pistil and large flowers? Finally, genomic sequencing of the populations will allow us to examine whether the specific genomic regions selected during this experiment overlap with those targeted during natural divergence in *M. lewisii*, directly testing the predictability and repeatability of adaptation to a shared pollinator.

## Acknowledgements

We would like to thank the Haverford undergraduate students enrolled in BIO301 Pollination Ecology Superlab in 2022, 2023, 2024, and 2026. In addition, we thank Emily Wilkins and the greenhouse crew for taking care of the plants. This work is supported by a NSF EDGE grant IOS: 2319722 and the start-up funds for F. Peng from Haverford College.

## Author contributions

F. P.: Conceptualization, Methodology, Funding acquisition, Supervision, Writing - original draft and review. CvV.: Data collection, Formal analysis. B. W. : Data collection, Validation. A. T. Data analysis. S. T. Data analysis. L. S.: Data collection

## Data Availability

Data will be available upon publication

